# Effects of contrast water therapy on performance, circulatory function, and fatigue in collegiate freestyle swimmers

**DOI:** 10.1101/2025.06.17.660270

**Authors:** Kazuki Kino, Mitsuo Neya, Yuya Watanabe, Noriyuki Kida

## Abstract

This study aimed to examine the effects of contrast water therapy (CWT) on recovery following high-intensity interval training in competitive collegiate swimmers. Fifteen male freestyle swimmers (mean age: 19.3 ± 1.1 years) participated in a crossover design, performing five 100-meter maximal-effort intervals under two conditions: CWT and passive rest (PAS). Each session included standardized warm-up, pre- and post-intervention measurements of blood pressure (BP), blood lactate concentration (LA), and subjective fatigue (FAS), followed by a second interval set. CWT consisted of 10 cycles of hot (40–41 °C, 60 s) and cold (20–21 °C, 30 s) full-body immersion. Performance metrics—swim time, stroke count, stroke length, and stroke velocity—were recorded during both interval sets. The results showed no significant differences between CWT and PAS in swim performance or blood pressure. However, a significant interaction was observed in stroke length during the first interval set, indicating a potential subtle benefit of CWT on swimming mechanics. Importantly, blood lactate concentrations were significantly lower after CWT compared to PAS (p < .001), and subjective fatigue was also reduced. These findings suggest that CWT promotes metabolic recovery, likely through enhanced peripheral circulation and lactate clearance, without negatively affecting cardiovascular parameters. Although CWT did not enhance immediate performance, its ability to reduce physiological and perceptual fatigue indicates its value in managing accumulated fatigue during training cycles. In conclusion, contrast water therapy appears to be a practical and effective recovery strategy for competitive swimmers, especially in supporting lactate clearance and reducing fatigue following high-intensity training. Future studies should explore optimal immersion protocols and assess long-term benefits across varied performance levels.

## Introduction

Competitive swimming is a sport in which athletes race a designated distance, typically ranging from 50 to 1500 meters, using specific strokes. The duration of physical exertion varies greatly by event—from as short as 20 seconds to over 15 minutes—resulting in a wide spectrum of physiological energy systems being utilized. Sprint events are primarily driven by anaerobic metabolism, whereas long-distance events rely on aerobic metabolism. Regardless of event type, competitive swimmers engage in prolonged, high-intensity training sessions over extended periods [1]. It has been reported that typical daily training distances range from 9,000 to 18,000 meters, involving significant physical strain [2]. Furthermore, the widespread availability of heated indoor pools has allowed for competitions to be held year-round, effectively lengthening both training and competitive seasons. These developments further emphasize the critical importance of recovery strategies to mitigate fatigue and maintain performance [3].

Given this context, the establishment of effective recovery strategies has become a key issue for maximizing performance and training efficiency in competitive swimmers. In recent years, various recovery methods have been developed in response to this growing need. Among them, water immersion-based recovery techniques have attracted attention as potential tools for reducing fatigue after training and competition [4, 5] . These methods leverage the properties of hydrostatic pressure, buoyancy, and thermal stimulation (hot/cold), which can promote improved circulation, reduce muscle inflammation, and alleviate subjective fatigue [6, 7]. Specifically, hydrostatic pressure enhances venous return and cardiac output, contributing to lactic acid removal and inflammation reduction [6]. Buoyancy, by reducing gravitational stress on the body, decreases musculoskeletal load and energy expenditure, helping to ease fatigue [7]. Additionally, alternating hot and cold stimuli can alter skin temperature, which may relieve muscle soreness and inflammation and promote relaxation through enhanced blood flow [8, 9].

Contrast water therapy (CWT), which alternates between hot and cold water immersion, is widely employed to promote blood flow and autonomic regulation, thereby facilitating rapid recovery [9]. Numerous studies have reported that CWT is effective in reducing muscle soreness, alleviating fatigue, and improving subjective recovery [10, 11]. However, the outcomes of CWT may vary depending on the sport, specific protocol, and individual characteristics, and consistent conclusions about its effectiveness remain elusive. In particular, research focused on competitive swimmers is limited, and comprehensive evaluations of CWT’s benefits in this population are scarce [4, 12]. While some studies suggest that CWT may contribute to lactate removal and fatigue reduction [10, 13, 14], its direct impact on performance remains inconclusive.

Swimmers engage in highly specific aquatic training for prolonged durations, distinguishing them from athletes in other sports. Therefore, recovery strategies should be examined with a swimmer- specific perspective. In practice, various recovery methods are used alongside CWT. For example, icing is commonly used to reduce localized muscle damage and inflammation [12] , while active recovery is believed to promote blood flow and facilitate metabolite clearance [3]. Massage and compression garments are also widely employed to reduce perceived fatigue and muscle soreness [13]. Compared to these methods, CWT uniquely combines hydrostatic pressure and thermal stimulation, potentially offering simultaneous benefits in circulation and inflammation control [7]. This makes it a practical and potentially effective method in applied settings.

Therefore, this study aimed to investigate the effects of contrast water therapy following high-intensity 100-meter training in collegiate male freestyle swimmers. Specifically, we evaluated its impact from multiple perspectives, including swimming performance, circulatory function, and subjective fatigue. By standardizing water temperature and immersion protocols, the present study seeks to enhance the practical applicability of CWT and contribute to the development of evidence-based recovery strategies in competitive swimming.

## Methods

### 2.1 Participants

The participants in this study were 15 male collegiate swimmers specializing in freestyle (mean age: 19.3 ± 1.1 years; height: 172.4 ± 4.5 cm; weight: 68.4 ± 5.9 kg). All swimmers belonged to teams competing in Japan’s collegiate second division and had an average of 11.1 ± 2.6 years of competitive swimming experience. The physical characteristics and personal best times of the participants are shown in Table 1.

**Table 1.** Physical characteristics of subiects.

| Variables | n=15 |
| --- | --- |
| Age (y) | 19.3 ± 1.12 |
| Height (cm) | 172.4 ± 4.52 |
| Weight (kg) | 68.4 ± 5.94 |
| BMI | 23.0 ± 1.44 |
| Body fat (%) | 12.1 ± 2.40 |
| LBM (kg) | 57.2 ± 4.04 |
| PB time (s) | 52.2 ± 1.23 |
| Career (y) | 11.1 ± 2.66 |

Values are represent as mean and SD
BMI, body mass index: LBM, lean body mass
PB, personal best in 100 m front crawl race

Prior to participation, all individuals were informed orally and in writing about the study’s purpose, procedures, and their right to withdraw at any time. Written informed consent was obtained from all participants. This study was approved by the Research Ethics Committee of the Faculty of Physical Education at Tenri University (Approval No. H29-017).

### 2.2 Physical Characteristics

Body weight, body fat percentage, fat-free mass, and lean body mass were measured using a multi-frequency body composition analyzer (MC-780A, TANITA Corporation).

### 2.3 Experimental Timeline

The timeline of the measurements is illustrated in Figure 1. A crossover design was adopted in which each participant underwent both contrast water therapy (CWT) and seated rest (PAS: passive rest) conditions on separate days.

**Fig 1.**
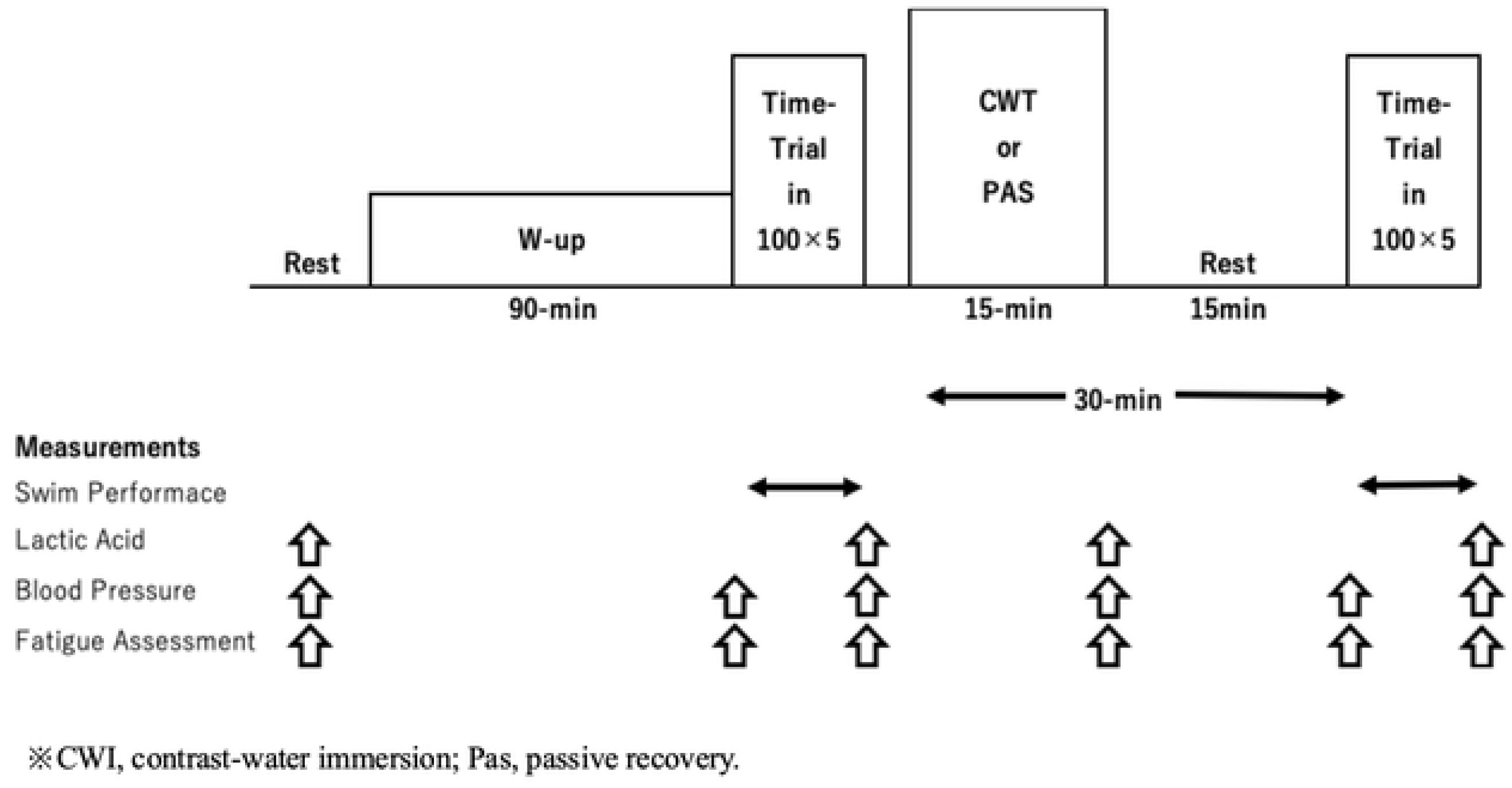
**Timeline of experimental protocol.**

Before the intervention, resting blood pressure (BP), blood lactate concentration (LA), and subjective fatigue (FAS: Fatigue Assessment Scale) were measured. Participants then completed a 90-minute pool-based warm-up that included swim, pull, and kick exercises totaling 1,000–2,000 meters.

Following the warm-up, participants performed five sets of 100-meter freestyle interval swims (work-to-rest ratio of 1:0.5), and were instructed verbally to maintain maximal effort throughout. Upon completion of the interval set, BP, LA, and FAS were measured again, followed by either CWT or seated rest.

Thirty minutes after the intervention, a second set of 100-meter interval swims was conducted. Measurements of BP, LA, and FAS were repeated after the second swim set. All measurements were performed in a pool facility maintained at an ambient temperature of 25–27 °C and humidity of 63– 68%. The swim trials were conducted in an indoor 25-meter pool with a water temperature maintained between 28.5–29.0 °C.

Previous studies (e.g., Parouty et al., 2010) have suggested that prolonged cold-water immersion may reduce core temperature and impair performance. Thus, the protocol in the present study was designed with these considerations in mind. Additionally, the temperature settings and immersion durations were based on reports indicating that ending with cold water may be more beneficial [7, 14]. Two indoor pools (approximately 1800 × 1200 mm) were used for hot and cold baths, placed 1 meter apart with transfer times kept under 5 seconds. Participants were seated and fully immersed up to their shoulders. Conversation during immersion was not permitted.

### 2.4 Swimming Performance, Circulatory Function, and Subjective Fatigue

A 60-fps camera was positioned at the center of the pool to film swimmers continuously. Swim time (ST), stroke count (SC), stroke velocity (SV), and stroke length (SL) were assessed during the 100-meter interval swims. Stroke count was defined as the number of single-arm strokes, and SL was calculated by dividing the swim distance by total stroke count.

Resting and post-training LA levels and pre- and post-intervention BP were measured. Resting BP was measured in a seated position using an automatic sphygmomanometer (HEM-720, OMRON), recording systolic (SBP) and diastolic blood pressure (DBP).

LA was assessed via fingertip capillary blood sampling using the Lactate Pro 2 analyzer (LT-1730, ARKRAY) immediately after BP measurement at rest and 3 minutes after training.

Subjective fatigue was assessed using a visual analog scale (VAS) as part of the Fatigue Assessment Scale (FAS), administered within 10 minutes after rest and training. The scale consisted of a 100-mm horizontal line with numerical indicators every 10 mm. The right end (100 mm) represented “the most severe fatigue ever experienced.”

### 2.5 Statistical Analysis

All data were expressed as mean ± standard deviation. Two-way repeated measures ANOVA was conducted to examine the effects of condition (CWT vs. PAS) and time point (see Figure 1) on swimming performance and physiological parameters. The assumption of sphericity was tested using Mauchly’s test; when violated, the Greenhouse-Geisser correction was applied, and adjusted degrees of freedom were used for the F-tests. Effect sizes were calculated using partial eta squared (ηp²). When significant interactions were found, Bonferroni-adjusted post hoc comparisons were performed for both within- and between-group comparisons. A significance level of p < .05 was used for all analyses. Statistical processing was conducted using IBM SPSS Statistics software (version 27).

## Results

### 3.1. Effects on Swimming Performance

The effects of the first 100-meter interval swim (TT1) on swim time (ST) are shown in Figure 2.

**Fig 2.**
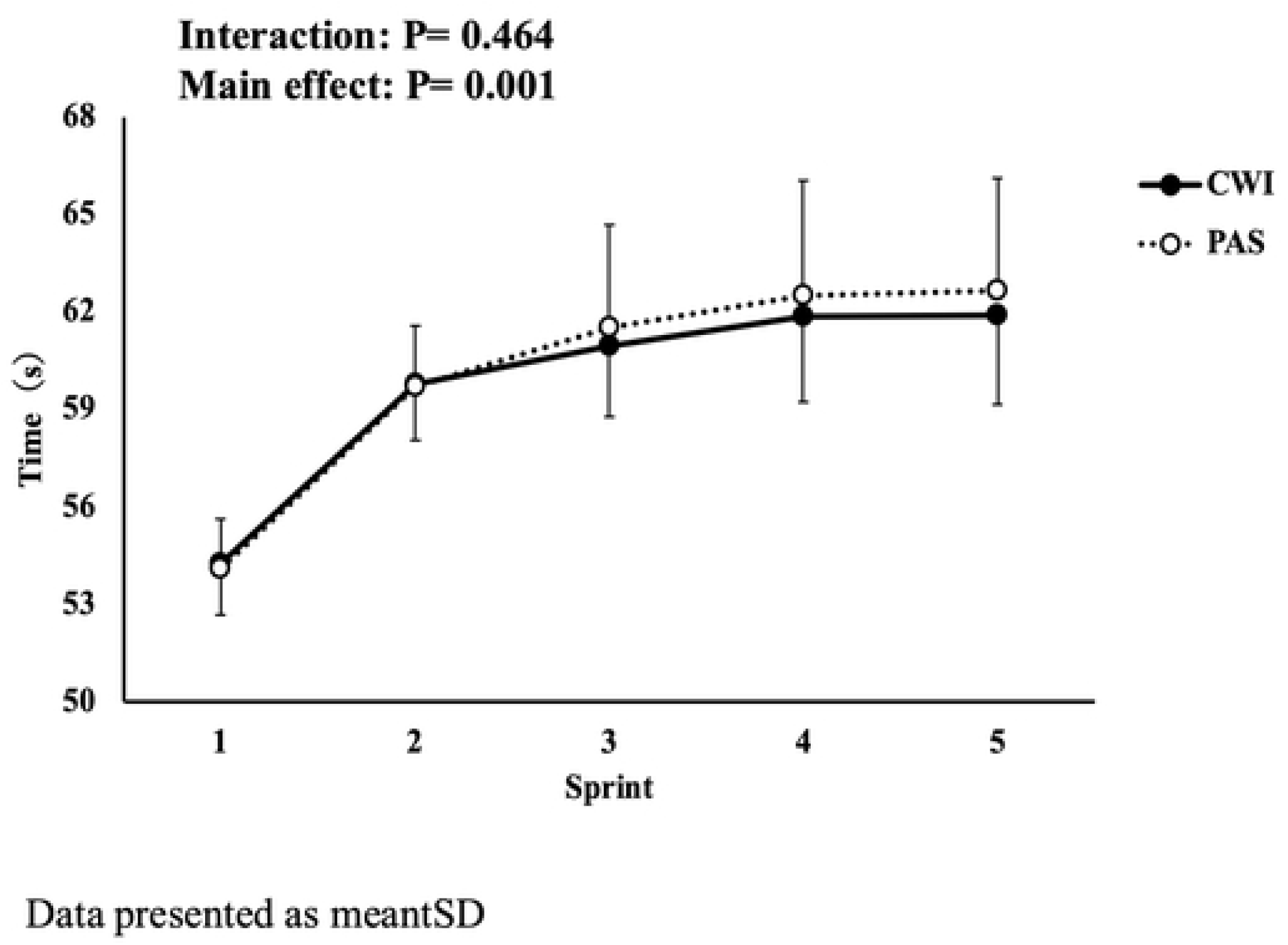
**Changes in the test time for the 100m freestyle crawl in Time Trial 1 for the control submersion (CWl) and passive recovery (PAS) groups.**

Two-way repeated measures ANOVA with Greenhouse-Geisser correction revealed no significant interaction between time and condition (F = 1.234, df = 34.857, p = 0.464, ηp² = 0.022).

Figure 3 shows the effects of the second 100-meter interval swim (TT2) on ST. Again, no significant interaction between time and condition was observed (F = 1.778, df = 49.778, p = 0.444, ηp² = 0.028).

**Fig 3.**
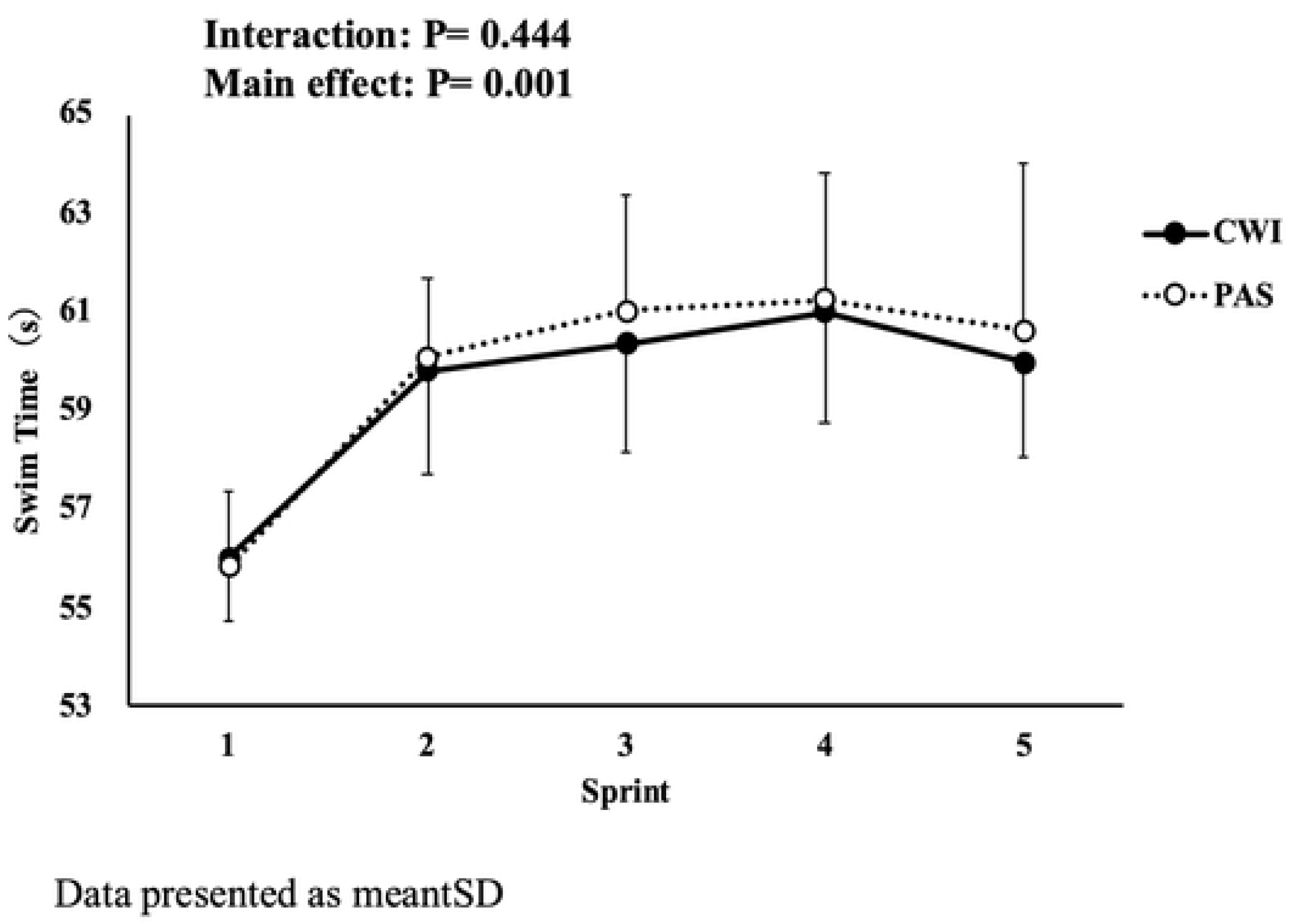
**Changes in the test time for the 100m freestyle crawl in Time Trial 2 for the control submersion (CWl) and passive recovery (PAS) groups.**

The effects of TT1 on stroke velocity (SV) was shown. No significant interaction was found (F = 1.267, df = 35.489, p = 0.487, ηp² = 0.017).

This shows the effects of TT2 on SV, which also did not reveal a significant interaction (F = 1.916, df = 53.638, p = 0.669, ηp² = 0.023).

The effects of TT1 on stroke length (SL) was shown. A significant interaction between time and condition was found (F = 1.806, df = 50.557, p < .005, ηp² = 0.201).

This shows the effects of TT2 on SL, which also did not reveal a significant interaction (F = 2.781, df = 77.857, p = 0.349, ηp² = 0.012).

The effects of TT1 on stroke count was shown. No significant interaction between time and condition was detected (F = 2.277, df = 63.749, p = 0.484, ηp² = 0.017).

Similarly, there was no significant interaction for the effect of TT2 on stroke count (F = 2.240, df = 62.726, p = 0.362, ηp² = 0.032).

### 3.2. Circulatory Function

Figure 4 shows the changes in systolic blood pressure (SBP) across measurement points, and

**Fig 4.**
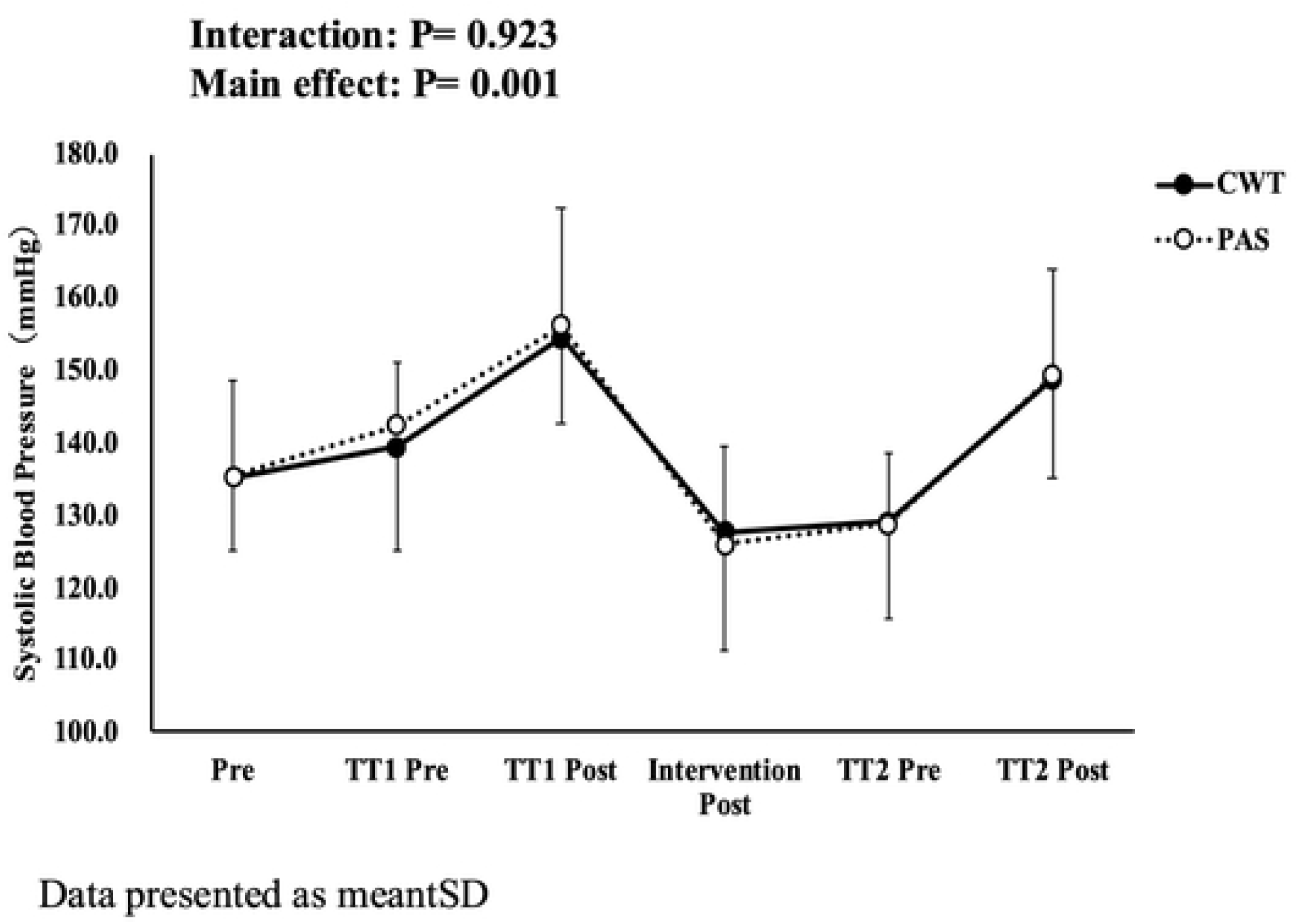
**Changes in systolic blood pressure in the control submersion (CWl) group and passive recovery (PAS) group at each measurement point.**

Figure 5 presents the changes in diastolic blood pressure (DBP). Figure 6 displays the blood lactate concentrations (LA). Two-way repeated measures ANOVA revealed no significant interaction between time and condition for both SBP and DBP.

**Fig 5.**
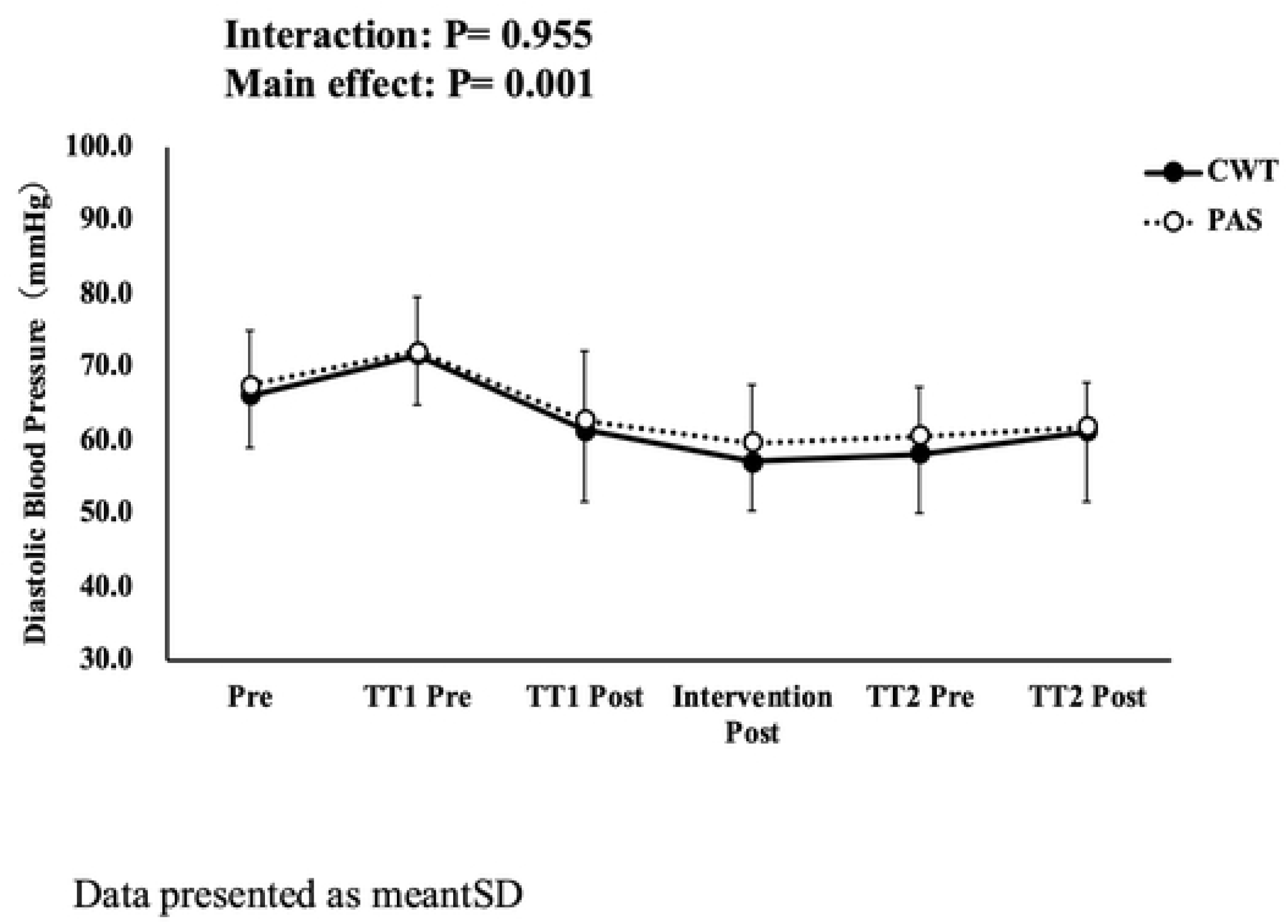
**Changes in diastolic blood pressure in the control submersion (CWl) group and passive recovery (PAS) group at each measurement point.**

**Fig 6.**
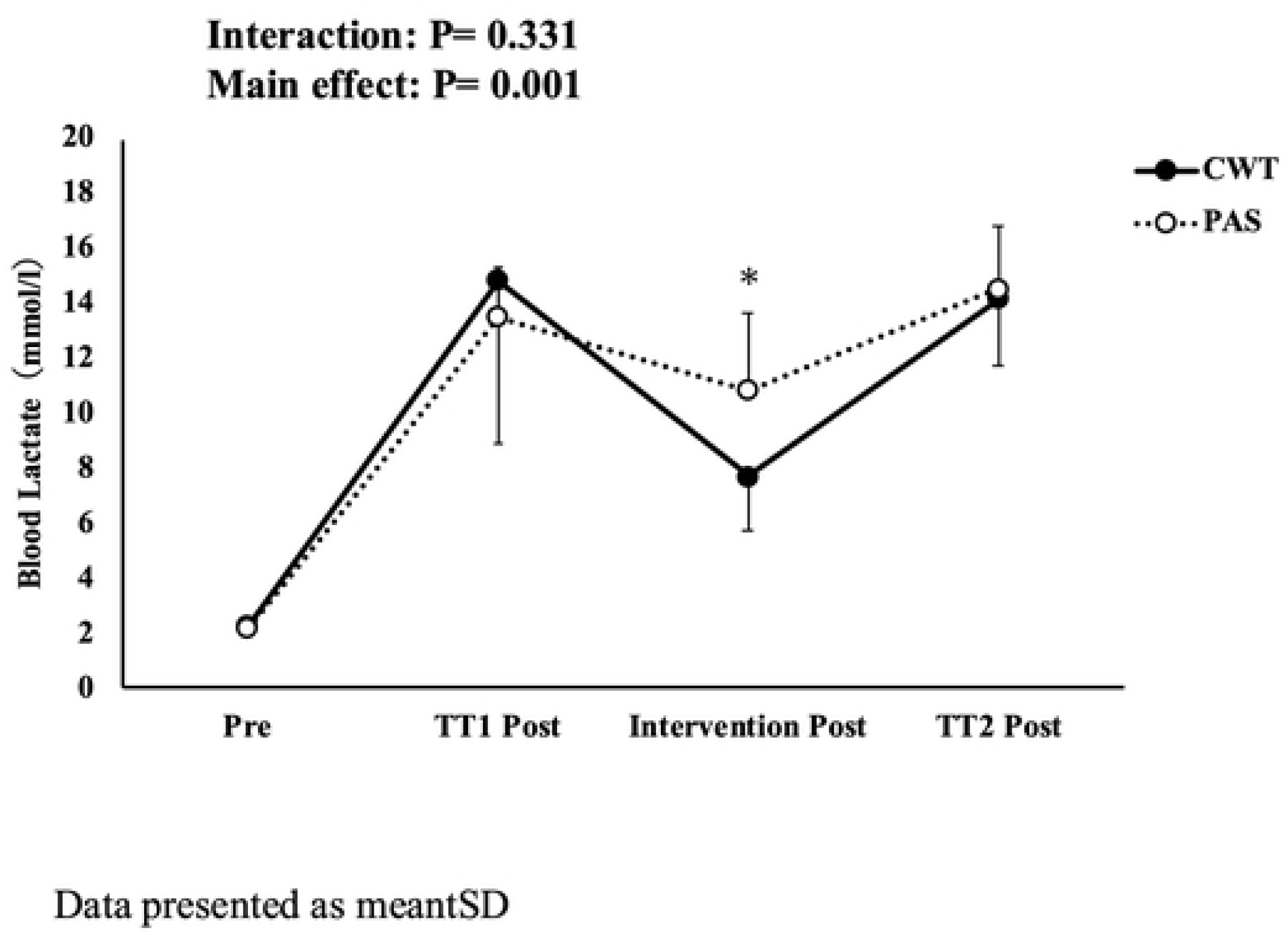
**Changes in blood lactate in the control submersion (CWl) group and passive recovery (PAS) group at each measurement point.**

However, a significant interaction was observed for LA (F = 3.84, p < .001, ηp² = 0.245), indicating that the changes in blood lactate concentrations differed between the contrast water therapy (CWT) and passive rest (PAS) conditions.

### 3.3. Subjective Fatigue

Figure 7 shows the effects of the intervention on subjective fatigue assessed by the Fatigue Assessment Scale (FAS). No significant interaction between time and condition was observed (F = 2.800, df = 78.405, p = 0.331, ηp² = 0.040). However, pairwise comparisons revealed a significant difference between pre- and post-intervention measurements (p < 0.021).

**Fig 7.**
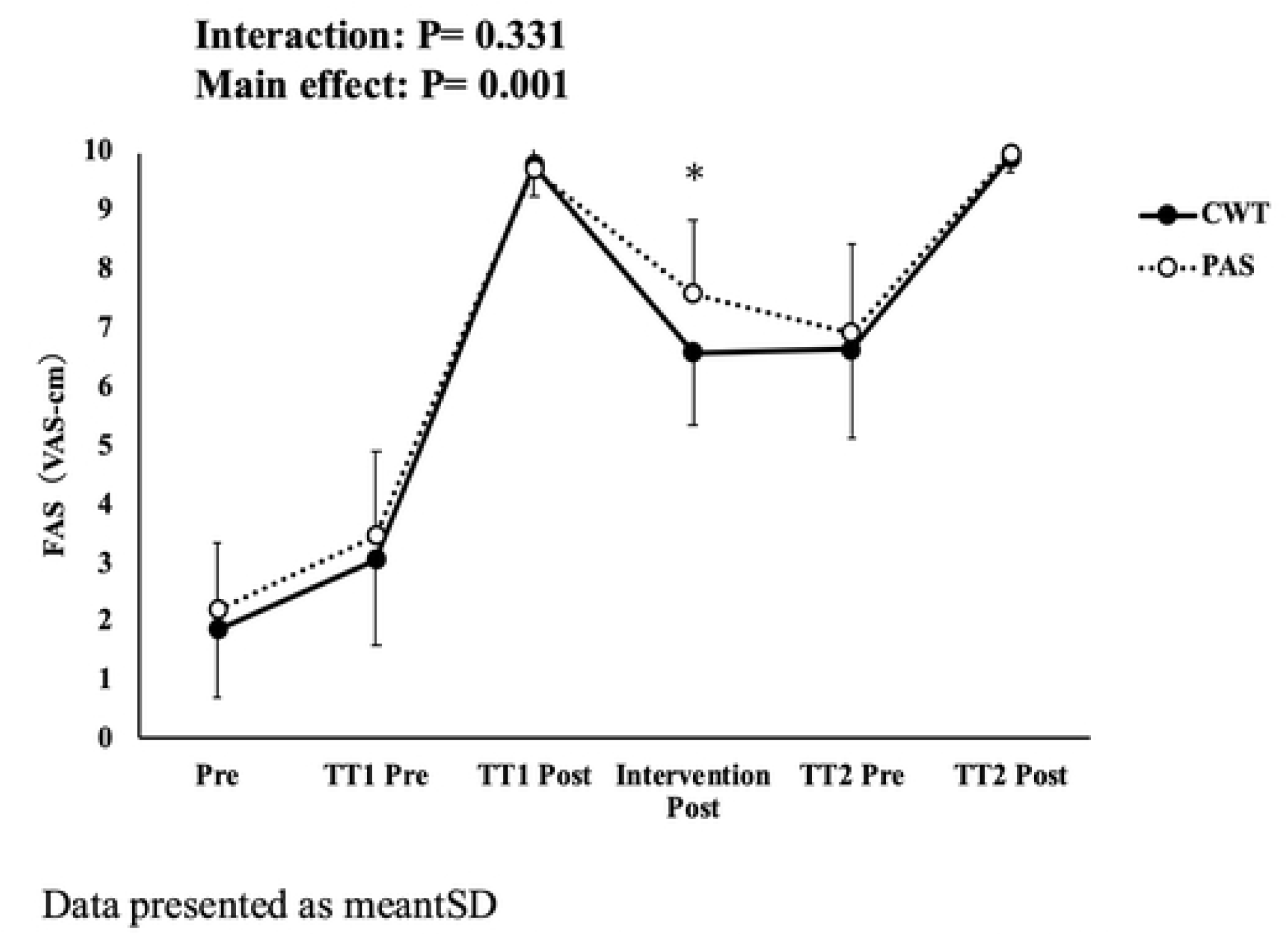
**Changes in FAS (VAS-cm) in the control submersion (CWl) group and passive recovery (PAS) group at each measurement point.**

## **4.** Discussion

### **A.** Swimming Performance

This study investigated the effects of contrast water therapy (CWT) on swimming performance, circulatory function, and subjective fatigue following high-intensity 100-meter training swims in collegiate swimmers. Participants underwent 15 minutes of alternating hot water immersion (40–41 °C, 60 seconds) and cold water immersion (20–21 °C, 30 seconds) across 10 cycles. The results showed that, compared to passive rest, CWT did not lead to significant differences in swim performance or blood pressure. However, CWT did facilitate blood lactate clearance and reduced subjective fatigue. In terms of swim performance, no significant differences were observed in swim time or most stroke-related variables during the 100-meter interval swims following CWT. However, while most stroke-related metrics showed no significant changes, a significant interaction in stroke length during TT1 suggests that certain aspects of swimming mechanics may be acutely influenced by CWT. This suggests that short-term CWT may not have an immediate effect on overall performance outcomes, although subtle mechanical changes could occur. These findings are consistent with those of [14], who reported that although six minutes of CWT effectively supported acute recovery in runners, it did not improve immediate performance.

Some studies have reported performance-enhancing effects of water-based recovery methods [15, 16]. [17] for instance, found that among elite swimmers, conducting CWT during recovery periods between 100-meter sprint swims led to improved repeated sprint performance. Although these results differ from the present study, it is important to note that most prior studies used single-effort maximal sprints powered predominantly by the ATP-CP and glycolytic systems. In contrast, the present study employed interval training, where energy supply capacity over multiple bouts may have influenced performance outcomes. This difference in energy system demands may partially explain the inconsistency in findings.

### **B.** Physiological Responses

This study found that contrast water therapy (CWT) promoted blood lactate clearance. This outcome suggests that the alternating thermal stimuli of CWT may increase peripheral blood flow and enhance lactate metabolism. During the high-intensity 100-meter interval swims in this study, the demand for rapid ATP production likely activated the glycolytic energy system. Under insufficient oxygen conditions, pyruvate is converted into lactate, resulting in elevated blood lactate concentrations [18]. Although traditional views associated elevated blood lactate with fatigue [18], recent studies have highlighted the greater influence of intramuscular pH on fatigue [19, 20]. Nevertheless, increased blood lactate levels have been linked to impaired muscle function and reduced exercise performance [21, 22]. [23] reported that, among elite male swimmers, high-intensity swim tests that raised lactate levels were associated with reduced stroke index. Furthermore, elevated blood lactate is thought to lower body fluid pH, inhibit glycolytic ATP production, impair muscle contraction, and delay physical recovery after exercise [24].

These findings underscore the importance of promoting lactate clearance as a component of effective recovery strategies. Future research should examine whether improved lactate clearance through CWT can influence performance over longer recovery periods.

CWT has been reported to increase peripheral blood flow [25] and intramuscular circulation [26]. Although the precise vascular responses to thermal stimulation remain unclear, several studies have indicated that alternating vasoconstriction and vasodilation during CWT can enhance lactate clearance [9, 25, 27, 28]. In the present study, post-intervention lactate concentrations were significantly lower in the CWT group compared to the passive rest group. This finding supports the hypothesis that CWT promotes recovery by facilitating lactate removal via enhanced blood flow.

However, it should be noted that some studies have suggested that the increased blood flow during CWT primarily affects the skin’s surface and may have limited influence on deep tissue circulation [29]. Therefore, further research is warranted to clarify the extent to which CWT affects lactate clearance, particularly in deeper muscle tissues.

Previous studies on post-exercise CWT have shown similar results. [30] found that active recovery promoted lactate clearance in 17 elite swimmers after maximal 200-meter freestyle efforts. Likewise, [31] reported that both active recovery and CWT significantly reduced blood lactate concentrations following a Wingate ergometer test in 14 collegiate hockey players. [9] also suggested that cold-water immersion at 10–15°C may be effective for lactate clearance.

Although the water temperature used for cold immersion in the present study (∼20°C) was slightly higher than in previous studies, conducting 10 cycles of CWT still resulted in improved lactate clearance. These findings suggest that CWT may be comparable to active recovery in terms of lactate removal, while potentially imposing a lower physical burden than cold water immersion alone. Thus, CWT appears to be a practical and effective recovery method in settings where rapid recovery is required.

In contrast, no significant changes were observed in systolic or diastolic blood pressure following CWT. [26] suggested that thermal stimulation during CWT may induce vascular responses leading to blood pressure changes. However, considering that the effects of CWT are largely limited to peripheral circulation, it is possible that no changes in blood pressure were detected in this study.

Future studies should explore the influence of various temperature settings and durations on lactate clearance. Additionally, the effects of CWT on deep tissue circulation should be assessed using more precise measurement techniques.

### **C.** Subjective Fatigue

This study demonstrated that contrast water therapy (CWT) was effective in reducing subjective fatigue. [6] reported that full-body immersion baths led to greater reductions in fatigue compared to showering. Similarly, [32] found that CWT performed after high-intensity training resulted in lower fatigue index scores.

The present findings suggest that the hydrostatic pressure, buoyancy, and thermal stimulation provided by full-body immersion during CWT may contribute to the reduction in perceived fatigue. These physical properties are known to promote relaxation, reduce musculoskeletal stress, and enhance circulation, thereby facilitating recovery and reducing the subjective sensation of fatigue.

## Practical Applications

The present study found that contrast water therapy (CWT) did not produce immediate effects on swimming performance or blood pressure. Therefore, its utility for enhancing performance directly after competition may be limited.

However, CWT was effective in promoting blood lactate clearance and reducing subjective fatigue, suggesting its potential value in medium-to long-term fatigue management and recovery support. In practical settings, individual variability and differences in response depending on water temperature and immersion duration must be taken into account. Accordingly, it is important to flexibly incorporate CWT in combination with other recovery methods.

Understanding the unique characteristics of CWT and applying it appropriately based on the athlete’s condition and recovery goals will be essential for maximizing its benefits.

## Conclusion

This study investigated the effects of contrast water therapy (CWT) following high-intensity 100-meter interval training in collegiate swimmers. While CWT did not produce immediate effects on swimming performance or blood pressure, it significantly promoted blood lactate clearance and reduced subjective fatigue.

These findings suggest that CWT is a useful recovery method that may a useful recovery method that may contribute to maintaining athletic performance over timeathletic performance by facilitating physiological and perceptual recovery. Further research is warranted to explore its effectiveness under a wider range of conditions and in athletes of varying performance levels, in order to establish optimal protocols for its use in competitive swimming.

## Acknowledgments

The authors would like to thank all participating students involved in this study. Special thanks are also extended to the staff and coaches for their valuable support.

## References

1. Maglischo EW. Swimming fastest: Human Kinetics; 2003. 744 p.

2. COSTILL DL. The 1985 C. H. McCloy Research Lecture Practical Problems in Exercise Physiology Research. RESEARCH QUARTERLY FOR EXERCISE AND SPORT. 1985;56:7. doi: 10.1080/02701367.1985.10605344.

3. Bishop PA, Jones E, Woods AK. Recovery from training: a brief review: brief review. J Strength Cond Res. 2008;22(3):1015–24. doi: 10.1519/jsc.0b013e31816eb518. PubMed PMID: 18438210.

4. Peeling P, Fulton S, Sim M, White J. Recovery effects of hyperoxic gas inhalation or contrast water immersion on the postexercise cytokine response, perceptual recovery, and next day exercise performance. J Strength Cond Res. 2012;26(4):968–75. doi: 10.1519/jsc.0b013e31822dcc5b. PubMed PMID: 22415455.

5. Parouty J, Al Haddad H, Quod M, Lepretre PM, Ahmaidi S, Buchheit M. Effect of cold water immersion on 100-m sprint performance in well-trained swimmers. Eur J Appl Physiol. 2010;109(3):483–90. Epub 20100217. doi: 10.1007/s00421-010-1381-2. PubMed PMID: 20162301.

6. Goto Y, Hayasaka S, Kurihara S, Nakamura Y. Physical and Mental Effects of Bathing: A Randomized Intervention Study. Evid Based Complement Alternat Med. 2018;2018:9521086. Epub 20180607. doi: 10.1155/2018/9521086. PubMed PMID: 29977318; PubMed Central PMCID: PMCPMC6011066.

7. Wilcock IM, Cronin JB, Hing WA. Physiological response to water immersion: a method for sport recovery? Sports Med. 2006;36(9):747–65. doi: 10.2165/00007256-200636090-00003. PubMed PMID: 16937951.

8. Higgins TR, Greene DA, Baker MK. Effects of Cold Water Immersion and Contrast Water Therapy for Recovery From Team Sport: A Systematic Review and Meta-analysis. J Strength Cond Res. 2017;31(5):1443–60. doi: 10.1519/jsc.0000000000001559. PubMed PMID: 27398915.

9. Vaile J, Halson S, Gill N, Dawson B. Effect of hydrotherapy on recovery from fatigue. Int J Sports Med. 2008;29(7):539–44. Epub 20071130. doi: 10.1055/s-2007-989267. PubMed PMID: 18058595.

10. Pelana R, Maulana A, Winata B, Widiastuti W, Sukur A, Kuswahyudi K, et al. Effect of contrast water therapy on blood lactate concentration after high-intensity interval training in elite futsal players. Physiotherapy Quarterly. 2019;27(3):12–9. doi: 10.5114/pq.2019.86463.

11. Versey NG, Halson SL, Dawson BT. Effect of contrast water therapy duration on recovery of running performance. Int J Sports Physiol Perform. 2012;7(2):130–40. Epub 20111212. doi: 10.1123/ijspp.7.2.130. PubMed PMID: 22173197.

12. Bleakley C, McDonough S, MacAuley D. The use of ice in the treatment of acute soft-tissue injury: a systematic review of randomized controlled trials. Am J Sports Med. 2004;32(1):251–61. doi: 10.1177/0363546503260757. PubMed PMID: 14754753.

13. Barnett A. Using recovery modalities between training sessions in elite athletes: does it help? Sports Med. 2006;36(9):781–96. doi: 10.2165/00007256-200636090-00005. PubMed PMID: 16937953.

14. Versey N, Halson S, Dawson B. Effect of contrast water therapy duration on recovery of cycling performance: a dose-response study. Eur J Appl Physiol. 2011;111(1):37–46. Epub 20100901. doi: 10.1007/s00421-010-1614-4. PubMed PMID: 20809231.

15. Buchheit M, Al Haddad H, Chivot A, Lepretre PM, Ahmaidi S, Laursen PB. Effect of in-versus out-of-water recovery on repeated swimming sprint performance. Eur J Appl Physiol. 2010;108(2):321–7. Epub 20091001. doi: 10.1007/s00421-009-1212-5. PubMed PMID: 19795131.

16. Fajar MK, Hariyanto A, Wahjuni ES, Hamzah SH, Wijono W, Sidik RM, et al. Cold water immersion as an effective recovery method: its impact on heart rate and lactate levels post exercise. Retos. 2024;61:440–7. doi: 10.47197/retos.v61.108794.

17. Zeinab Rezaee FEaSMM. Which Temperature During the Water Immersion Recovery Is the Best after a Sprint Swimming? World Applied Sciences Journal. 2012:1403–8.

18. Gladden LB. Lactate metabolism: a new paradigm for the third millennium. J Physiol. 2004;558(Pt 1):5–30. Epub 20040506. doi: 10.1007/s00421-009-1212-5. PubMed PMID: 15131240; PubMed Central PMCID: PMCPMC1664920.

19. Bangsbo J, Graham T, Johansen L, Saltin B. Muscle lactate metabolism in recovery from intense exhaustive exercise: impact of light exercise. J Appl Physiol (1985). 1994;77(4):1890-5. doi: 10.1152/jappl.1994.77.4.1890. PubMed PMID: 7836214.

20. Toubekis AG, Douda HT, Tokmakidis SP. Influence of different rest intervals during active or passive recovery on repeated sprint swimming performance. Eur J Appl Physiol. 2005;93(5-6):694–700. Epub 20041120. doi: 10.1007/s00421-004-1244-9. PubMed PMID: 15778899.

21. Hogan MC, Gladden LB, Kurdak SS, Poole DC. Increased [lactate] in working dog muscle reduces tension development independent of pH. Med Sci Sports Exerc. 1995;27(3):371–7. PubMed PMID: 7752864.

22. Westerblad H, Allen DG. Changes of intracellular pH due to repetitive stimulation of single fibres from mouse skeletal muscle. J Physiol. 1992;449:49–71. doi: 10.1113/jphysiol.1992.sp019074. PubMed PMID: 1522520; PubMed Central PMCID: PMCPMC1176067.

23. Aujouannet YA, Bonifazi M, Hintzy F, Vuillerme N, Rouard AH. Effects of a high-intensity swim test on kinematic parameters in high-level athletes. Appl Physiol Nutr Metab. 2006;31(2):150–8. doi: 10.1139/h05-012. PubMed PMID: 16604133.

24. Huang T, Liang Z, Wang K, Miao X, Zheng L. Novel insights into athlete physical recovery concerning lactate metabolism, lactate clearance and fatigue monitoring: A comprehensive review. Front Physiol. 2025;16:1459717. Epub 20250325. doi: 10.3389/fphys.2025.1459717. PubMed PMID: 40200988; PubMed Central PMCID: PMCPMC11975961.

25. Fiscus KA, Kaminski TW, Powers ME. Changes in lower-leg blood flow during warm-, cold-, and contrast-water therapy. Arch Phys Med Rehabil. 2005;86(7):1404–10. doi: 10.1016/j.apmr.2004.11.046. PubMed PMID: 16003672.

26. Shadgan B, Pakravan AH, Hoens A, Reid WD. Contrast Baths, Intramuscular Hemodynamics, and Oxygenation as Monitored by Near-Infrared Spectroscopy. J Athl Train. 2018;53(8):782–7. Epub 20180913. doi: 10.4085/1062-6050-127-17. PubMed PMID: 30212235; PubMed Central PMCID: PMCPMC6188085.

27. Hing WA, White SG, Bouaaphone A, Lee P. Contrast therapy--a systematic review. Phys Ther Sport. 2008;9(3):148–61. Epub 20080722. doi: 10.1016/j.ptsp.2008.06.001. PubMed PMID: 19083715.

28. Ingram J, Dawson B, Goodman C, Wallman K, Beilby J. Effect of water immersion methods on post-exercise recovery from simulated team sport exercise. J Sci Med Sport. 2009;12(3):417–21. Epub 20080611. doi: 10.1016/j.jsams.2007.12.011. PubMed PMID: 18547863.

29. Myrer JW, Draper DO, Durrant E. Contrast therapy and intramuscular temperature in the human leg. J Athl Train. 1994;29(4):318–22. PubMed PMID: 16558294; PubMed Central PMCID: PMCPMC1317806.

30. Ali Rasooli S, Koushkie Jahromi M, Asadmanesh A, Salesi M. Influence of massage, active and passive recovery on swimming performance and blood lactate. J Sports Med Phys Fitness. 2012;52(2):122–7. PubMed PMID: 22525646.

31. MARK G. SAYERS AMC, & JO G. SANDERS. Effect of whole-body contrast-water therapy on recovery from intense exercise of short duration. European Journal of Sport Science. 2011;11(4):293–302. doi: 10.1080/17461391.2010.512365.

32. Li T, Xiang H, Li L, Zhao C. Effects of contrast water therapy on proprioception of the knee joint and degree of fatigue in sprinters after high intensity training. Am J Transl Res. 2024;16(6):2492–500. Epub 20240615. doi: 10.62347/vgsh1115. PubMed PMID: 39006297; PubMed Central PMCID: PMCPMC11236656.

